# Real-time monitoring of drug pharmacokinetics within tumor tissue in live animals

**DOI:** 10.1101/2021.07.03.451023

**Authors:** Ji-Won Seo, Kaiyu Fu, Santiago Correa, Michael Eisenstein, Eric A. Appel, H. Tom Soh

## Abstract

The efficacy and safety of a chemotherapy regimen fundamentally depends on its pharmacokinetics. This is currently measured based on blood samples, but the abnormal vasculature and physiological heterogeneity of the tumor microenvironment can produce radically different drug pharmacokinetics relative to the systemic circulation. We have developed an implantable microelectrode array sensor that can collect such tissue-based pharmacokinetic data by simultaneously measuring intratumoral pharmacokinetics from multiple sites. We employ gold nanoporous microelectrodes that maintain robust sensor performance even after repeated tissue implantation and extended exposure to the tumor microenvironment. We demonstrate continuous *in vivo* monitoring of concentrations of the chemotherapy drug doxorubicin at multiple tumor sites in a rodent model, and demonstrate clear differences in pharmacokinetics relative to the circulation that could meaningfully affect drug efficacy and safety. This platform could prove valuable for preclinical *in vivo* characterization of cancer therapeutics, and may offer a foundation for future clinical applications.

## Introduction

Inter-individual differences in pharmacokinetics (PK) can profoundly affect the efficacy of drug treatment, particularly in the context of chemotherapy for cancer.^1, 2^ This variability makes it challenging to identify the appropriate therapeutic window for a given drug regimen. Underdosing reduces the likelihood of successful treatment, whereas overdosing can inflict severe damage on the kidney, liver, heart, and other organs.^3–7^ Currently, such PK data are typically collected via blood-based measurements of circulating drug concentrations, and a number of groups have even demonstrated the feasibility of real-time drug monitoring through the use of miniaturized implantable sensors.^8–11^ However, such circulation-based measurements do not adequately reflect drug absorption within the tumor tissue itself.^12–16^ This is because the microenvironment within tumors is complex, with unpredictable vascular permeability, heterogeneous and high interstitial fluid pressure, high cell density, and disorganized lymphatic drainage.^17–19^ These factors can collectively impede the continuous and homogenous penetration of drugs from plasma into the tumor tissue, resulting in notable differences in drug concentration between the plasma and different regions of the tumor. As such, measurements of drug concentrations in plasma can yield inaccurate assessments of PK, resulting in low therapeutic efficacy.

Currently, the only way to obtain tumor-specific PK measurements is through tissue specimens collected via needle-based biopsies. However, it is difficult to extrapolate overall drug penetration in the heterogeneous tumor tissue environment from a single sampling site, and this in turn leads to inaccurate PK assessment. Multiple biopsies would offer a more complete picture of tumor PK, but it is impractical to perform this invasive procedure repeatedly, because it is costly, time consuming and carries the risk of severe side effects—including tumor seeding.^20–23^ Even in animal models, multiple biopsies of tumors are technically challenging, and so researchers typically carry out PK studies by collecting samples from multiple animals sacrificed at different time-points, producing averaged population data that do not accurately capture intra-tumor variability from individual animals.^24^ Furthermore, these experiments are being performed *ex vivo*, and may not accurately capture the physiological behavior of a tumor within its *in vivo* milieu. More generally, such analyses are challenging to perform for a variety of reasons—for instance, the probe needs to withstand insertion into solid tissue with minimal sensor damage, and must be sufficiently resistant to biofouling to enable robust measurement over extended periods of time. Accordingly, there remains an unmet need for analytical tools that are capable of efficiently collecting accurate PK data from multiple tumor sites simultaneously.

Here, we demonstrate an electrochemical aptamer-based biosensor that enables robust, real-time, multi-site drug monitoring within tumor tissue in live animals. Our biosensor features a number of technical and design advances that enable it to overcome key limitations that have hindered past efforts to achieve drug monitoring within solid tissues. First, we make use of nanoporous gold microelectrodes that successfully minimize both the effects of fouling from biological matrices and the risk of sensor damage from tissue insertion. Second, each sensor incorporates several such microelectrodes so that we can monitor drug concentrations with a higher signal-to-noise ratio (SNR) at multiple sites within tumor tissue simultaneously. Finally, our sensor is flexible, offering a better match to the physical properties of surrounding tissue and thereby minimizing damage at the site of insertion. As a demonstration, we show that our microelectrode array sensor can monitor concentrations of the chemotherapy drug doxorubicin (DOX) at multiple positions in tumor tissue simultaneously. This enables us to collect *in situ* tumor-specific PK data that account for tissue heterogeneity within a single animal, revealing patterns of DOX distribution within the tumor tissue that differ starkly from those measured via the systemic circulation. These differences indicate that the latter metrics might prove misleading in the selection of an appropriate drug dose, and highlight the importance of *in situ* PK monitoring in the context of cancer therapeutics research. Our data indicate that our biosensor platform could offer a simple and robust tool for obtaining more physiologically relevant insights into drug PK and understanding the *in vivo* behavior of experimental drugs.

## Results and discussion

### Overview and fabrication of the sensor

Our sensor comprises an array of gold nanoporous microelectrodes integrated into a flexible, polyimide polymer-based probe, which can be implanted directly into tumor tissue in a live mouse (**Figure 1A**). This sensor is designed to collect temporal drug concentration profiles at multiple sites within tumor tissue simultaneously via multiple microelectrodes in real-time (**Figure 1B**). Based on the resulting measurements, the PK of the drug can then be assessed in terms of its absorption and elimination phases, where the former refers to the drug’s uptake into the tissue from the circulatory system and the latter describes the drug’s subsequent clearance from the tissue due to lymphatic drainage and other physiological processes.^25, 26^ The mean intratumoral PK of the drug—as calculated based on concentration data obtained at three different microelectrode channels—can then be compared to the systemic PK (**Figure 1C**).

**Figure 1.**
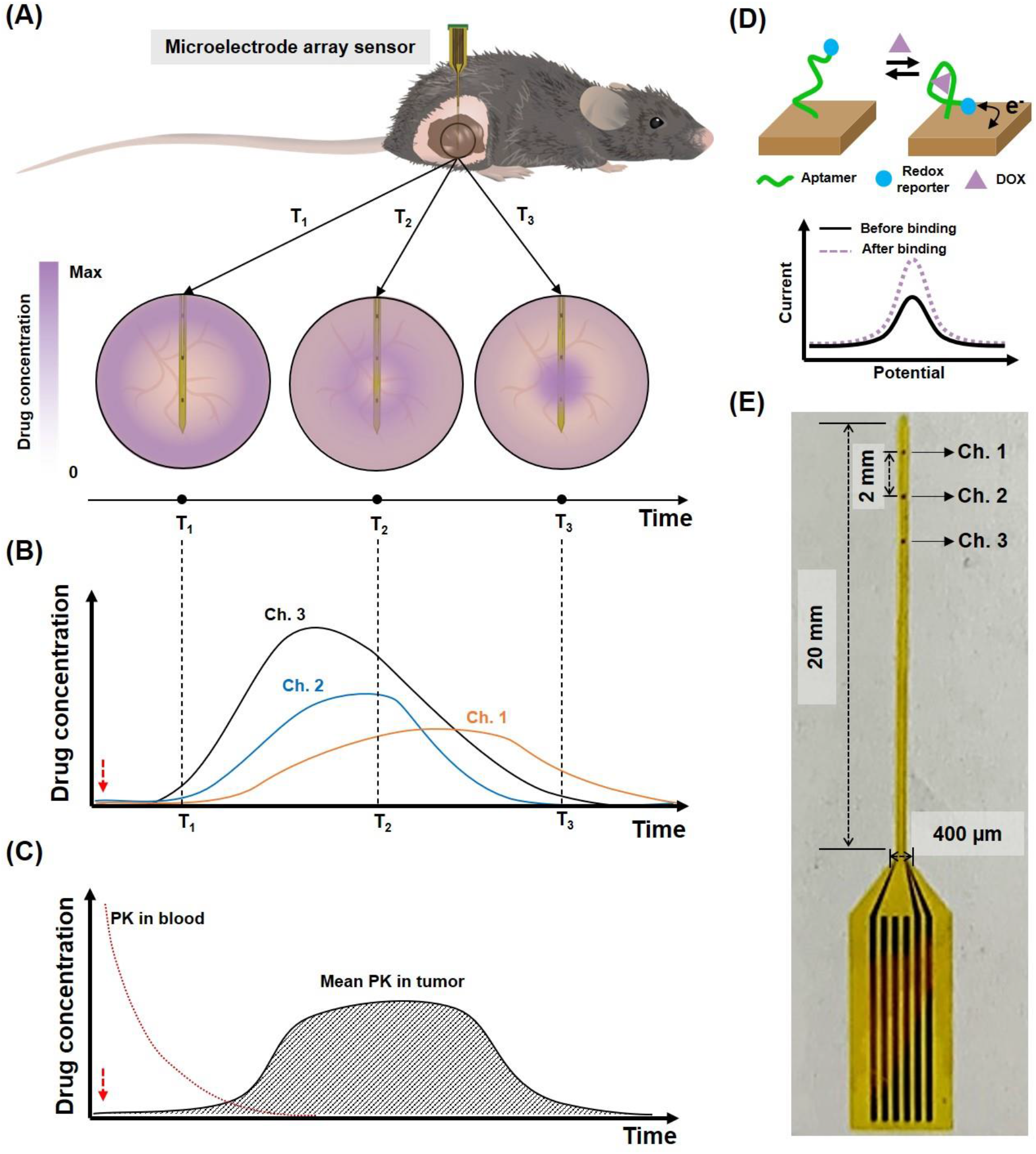
Gold nanoporous microelectrode array-based implantable electrochemical aptamer sensor. (A) Illustration of sensor implantation into tumor tissue in a live mouse (top) and detecting drug concentrations within the tumor at different time-points (T1, T2, and T3; bottom). (B) Illustration of real-time measurement of different drug concentrations at each microelectrode channel. Dotted lines represent the time-points from (A). (C) Illustration of averaged real-time drug concentrations from the three sensor channels versus drug concentrations from blood. Red dotted arrows in (B) and (C) indicate the time of drug injection. (D) Schematic of the aptamer-based drug detection mechanism. (E) Photo of our sensor.

Detection is achieved by functionalizing the surface of these gold microelectrodes with aptamers for the drug of interest that undergo a conformational change upon binding to their target. For the present work, we used a well-characterized aptamer that can bind DOX—a widely used chemotherapeutic drug.^27–31^ As described in our previous work^32^, the distal end of the aptamer is tagged with a methylene blue (MB) redox reporter; in the presence of the target, the aptamer undergoes a conformational change that increases the rate of electron transfer between the MB reporter and the electrode surface, thereby yielding an increase in current (**Figure 1D**). We applied square-wave voltammetry (SWV) to determine the signal gain, which is the ratio of current measurements from before and after target addition. Since the aptamer can reversibly bind and release its target, our sensor can continuously measure target concentration and kinetic information in real-time.

We recently reported that nanoporous structured electrodes offer greatly improved electrochemical detection sensitivity relative to planar electrodes, with a higher signal-to-noise ratio (SNR) due to reduced charge-screening effects.^33^ This enhanced sensitivity is important in this context, because *in vivo* experiments intrinsically have high background noise. The use of nanoporous microelectrodes should also facilitate long-term monitoring in tumor tissue, because aptamers residing within the nanopores are better protected against both mechanical damage during insertion and biofouling during implantation.^34, 35^ Each sensor comprises an array of several such nanoporous gold microelectrodes fabricated onto a flexible polyimide substrate, which forms a 400-μm x 20-mm shank with a thickness of 15 μm (**Figure 1E** and **S1**). The entire fabrication process is detailed in the Supporting Information (**Figure S2**). Briefly, an Au:Ag alloy film was deposited onto the polyimide substrate by co-sputtering of Au and Ag, after which the silver was dissolved in 69% nitric acid (**Figure S3A**). To prevent degradation of the polyimide substrate during this process, we added a gold bottom-protective layer before deposition of the Au:Ag alloy (**Figure S3B**). We then functionalized these microelectrodes with the MB-tagged DOX aptamer. For this work, we used a three-channel array consisting of 100 x 100 μm^2^ microelectrodes positioned with a 2-mm pitch. This pitch enables the sensor to measure a large area of tumor tissue simultaneously, while the relatively small area of the microelectrodes confers high spatial resolution compared to needle biopsies or metal wire-based sensors. The sensor was connected to a printed circuit board (PCB), which was in turn connected to a commercial potentiostat (**Figure S4**). The working electrode array in the sensor and a conventional Ag/AgCl reference electrode were used together for all measurements. After recording the SWV curves with the potentiostat, we utilized a custom MATLAB script to calculate the DOX concentration.

The mechanical flexibility conferred by our polyimide substrate is another essential feature of our *in vivo* sensor. It is well known that sensors made of silicon or metal wires lead to a mismatch in mechanical properties between the sensor (stiffness ~1 mN·m) and surrounding tissue (~100 nN·m)^36^, inflicting damage on the tumor tissue. Such tissue damage could lead to inaccurate measurement, prevent long-term monitoring, or cause additional tumor seeding.^22^ Our flexible probe minimizes such risks because its stiffness is sufficiently low (~270 nNom) to approach that of tumor tissue.^37–39^

### Characterizing sensor performance of our microelectrode array

We initially tested the performance of our sensors in a series of *in vitro* experiments. We first confirmed that each microelectrode in our array has equivalent sensitivity. Briefly, we immersed our sensor in 1x saline sodium citrate (SSC) buffer and introduced 10 μM DOX after allowing the baseline signal to stabilize for 12 minutes. All three channels showed a similar response, producing an average ~53.9% signal gain with 1.4% variance between channels (**Figure 2A**). At t = 17 min, we washed the sensor, and the signal returned to baseline within minutes, confirming the capacity for continuous, real-time sensing. We next exposed our multi-channel sensor to increasing DOX concentrations, and observed a clear and proportional signal gain; our three electrodes exhibited a steady increase in signal gain from 2.9% at 500 nM to 78.1% at 30 μM, with just 3% variance in signal gain at each concentration across the three electrodes (**Figure 2B**). We calculated the average signal gain of each DOX concentration from the data obtained at the three electrodes, and then performed curve fitting for this average signal gain data after plotting versus DOX concentration (**Figure S5**). We used this curve for the calibration of DOX concentration from the signal gain in all *ex vivo* and *in vivo* measurements described throughout the manuscript.

**Figure 2.**
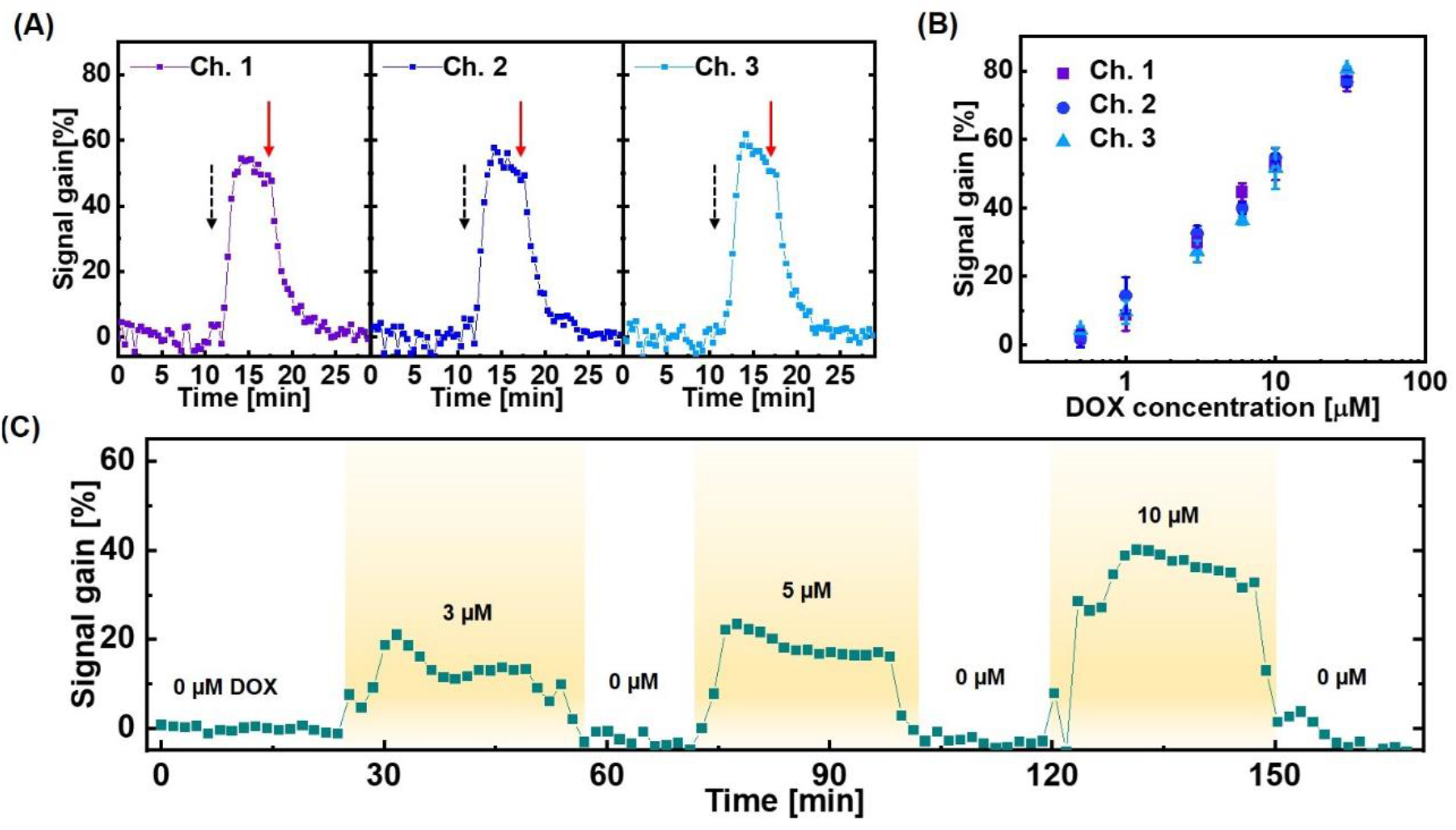
Assessing the consistency and reproducibility of microelectrode array measurements. (A) Continuous measurements of signal gain from each of our three channels at baseline, after adding a 10 μM bolus of DOX at t = 12 min (black dotted arrow), and after washing in 1x SSC buffer at t = 17 min (red solid arrow). (B) Signal gain from all three microelectrode channels at increasing DOX concentrations. Error bars were calculated from five different data points at each concentration. (C) Continuous measurements of signal gain in flowing fetal bovine serum containing different concentrations of DOX. SWV frequency = 200 Hz, amplitude = 50 mV. The data represent average signal gain from the continuously-measured signal gain data collected at the three channels.

We next set out to characterize the performance of our sensor in fetal bovine serum. We positioned our microelectrode array sensor vertically within a PDMS chamber, through which we continuously flowed undiluted serum with a peristaltic pump (**Figure S6**). We increased the DOX concentration in serum to 3 μM, 5 μM, or 10 μM at different time-points, and maintained each condition for 30 min. The aptamer-functionalized microelectrodes clearly responded to each DOX concentration, producing signal gains ranging from 13.2% at 3 μM DOX to 38.8% at 10 μM DOX (**Figure 2C**). Importantly, the sensor consistently returned to baseline when DOX was no longer present in serum, even after nearly three hours of continuous data collection—a time-scale that is standard for clinical DOX administration.

Biofouling of electrode surfaces poses a major problem for *in vivo* detection, and the adsorption of proteins present in the blood around and inside tumor tissue can render sensors unusable after a short period. Previous aptamer-based sensors have used a passivation layer on the electrode surface to mitigate this problem, but even with such measures, most aptamer-based sensors to date have reported a functional lifetime of no more than 12 hours in flowing blood.^40–42^ Based on prior work with nanostructured gold microelectrodes^34, 35^, we anticipated that our sensor would enable stable, long-term monitoring due to sequestration of the aptamers within nanopores, minimizing the effects of biofouling. To verify this, we measured the baseline signal of both planar and nanoporous microelectrodes in flowing serum for 16 h. As expected, the signal from planar microelectrodes worsened over time, and decreased to 10% of the baseline signal by 16 h, indicating severe biofouling and/or degradation of the aptamer (**Figure 3A**). In contrast, the nanoporous microelectrodes maintained 70% of the baseline signal after 16 h, indicating much more stable sensor performance in conditions that are highly conducive to biofouling. We next assessed the signal gain produced in response to a 3 μM DOX spike in serum—a standard clinical dose—at time zero versus after 16 h exposure to serum. At initial exposure, the nanoporous microelectrode produced an average signal gain of 11.1% versus 7.1% for the planar microelectrodes (**Figure 3B** and **S7**). After 16 h, the signal gain from the nanoporous electrodes decreased only slightly to 10.1% at 3 μM DOX, whereas the planar electrode sensor no longer produced a measurable signal gain. These results strongly suggest that the nanoporous gold microelectrodes are far less susceptible to biofouling and/or degradation of the DOX aptamer, and therefore better suited for long-term *in vivo* measurements.

**Figure 3.**
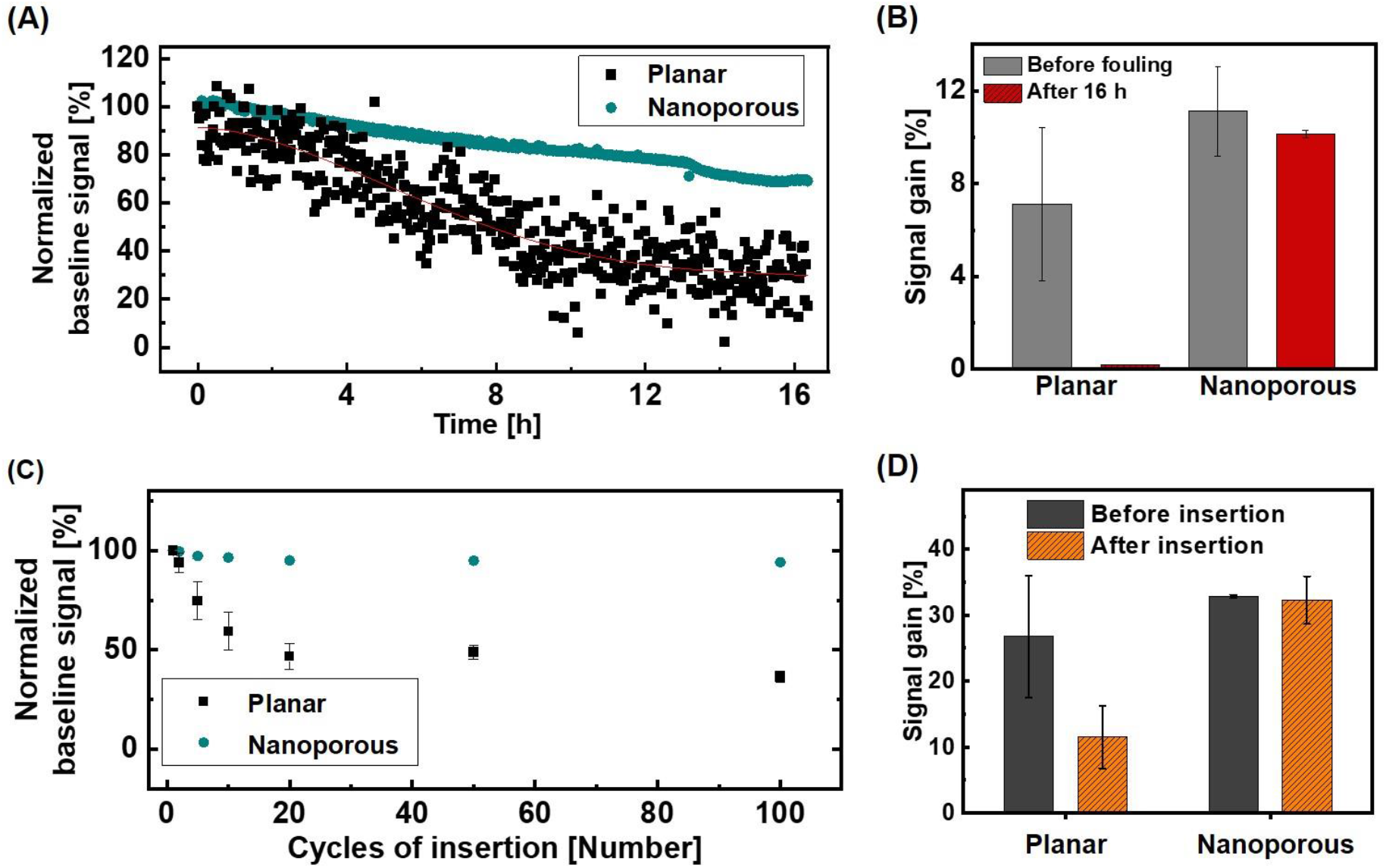
Stability of sensor performance in serum and tissue. (A) Representative data from a single channel showing baseline signal on gold nanoporous and planar microelectrodes over the course of 16 h in undiluted fetal bovine serum. (B) Average signal gain (n = 2 channels) produced by gold nanoporous and planar microelectrodes in the presence of 3 μM DOX in serum upon initial exposure (gray) and after 16 h in serum (red). (C) Averaged baseline signal change (n = 3 channels) for gold nanoporous and planar microelectrodes after multiple cycles of insertion into murine melanoma tumor tissue. (D) Averaged signal gain (n = 3 channels) from 6 μM DOX in 1x SSC on planar and nanoporous microelectrodes before initial insertion into tumor tissue and after 100 insertion cycles. SWV frequency = 200 Hz, Amplitude = 50 mV.

### Measurements from gold nanoporous microelectrodes inserted into tissue

We also anticipated that our nanoporous structured electrodes would offer protection against mechanical damage to the sensor surface, which could otherwise result in unstable and reduced signal after insertion. For comparative purposes, we fabricated three-channel planar microelectrode array sensors that were otherwise identical to our nanoporous microelectrode array sensors. We inserted planar and nanoporous gold microelectrode array sensors into tumor tissue and analyzed changes in the baseline signal over the course of multiple cycles of insertion. For these experiments, we used tumor tissue derived from the murine B16-F10 melanoma model, which was grown subcutaneously in the hind flank of a C57BL/6 mouse.^43^ In each insertion cycle, the sensor was inserted into tumor tissue and withdrawn immediately. The signal was then measured in 1x SSC buffer after each cycle of insertion. After 10 cycles, the average signal from planar microelectrodes decreased to 59.4% of the original baseline signal, and after 100 cycles, this signal was reduced to 36.4% of baseline (**Figure 3C**). In contrast, our nanoporous microelectrodes retained 94.1% of their baseline signal even after 100 cycles of insertion. These results clearly demonstrate that the gold nanoporous microelectrodes are very well suited for electrochemical detection in the context of solid tissue, with minimal mechanical damage after insertion to either the substrate or the functionalized aptamers.

We further confirmed this result by comparing the signal gain in response to 6 μM DOX in 1x SSC for these various microelectrodes before insertion into tumor tissue and after 100 cycles of insertion (**Figure 3D**). The planar and nanoporous microelectrodes respectively showed comparable average signal gain of 26.8% and 32.8% before insertion. But after 100 cycles, the signal gain from the planar microelectrodes sharply decreased to 11.5%, whereas the nanoporous electrode still maintained a 32.3% signal gain. We also assessed how well our probe performed in the context of tissues with different mechanical properties, and found that our nanoporous microelectrode sensor retains its baseline signal independent of the elastic modulus of the tissue environment (**Figure S8**), confirming the broad mechanical compatibility of our sensor design.

### *Ex vivo* real-time monitoring of DOX

We next assessed the real-time, multi-channel DOX detection capabilities of our sensor in surgically-removed tumor tissue (**Figure 4A**). We extracted ~100 mm^2^ of B16-F10 tumor tissue grown in a C57BL/6 mouse. After transferring the tumor tissue to a PDMS chamber within a small volume of buffer, we implanted our sensor into the middle of the tumor tissue such that all three channels of the array were embedded, with channel 1 closest to the center of the tissue. We then injected a 5 μg/g bolus of DOX adjacent to channel 3 and measured the DOX concentration at the three microelectrode channels (**Figure 4B**). All three clearly responded to this DOX spike in real-time, but each channel detected a different concentration. Channel 3, which was closest to the injection site, detected an average ~2.4 μM DOX, whereas the more distal channels 1 and 2 detected much lower average concentrations of ~0.8 and 0.7 μM, respectively. We next injected 500 μl 1x SSC buffer three times into the tumor to wash the drug out completely, and this treatment returned the measured concentration to zero. These results demonstrate that our system can discriminate spatial differences in the drug concentration profile at different positions within the tumor tissue.

**Figure 4.**
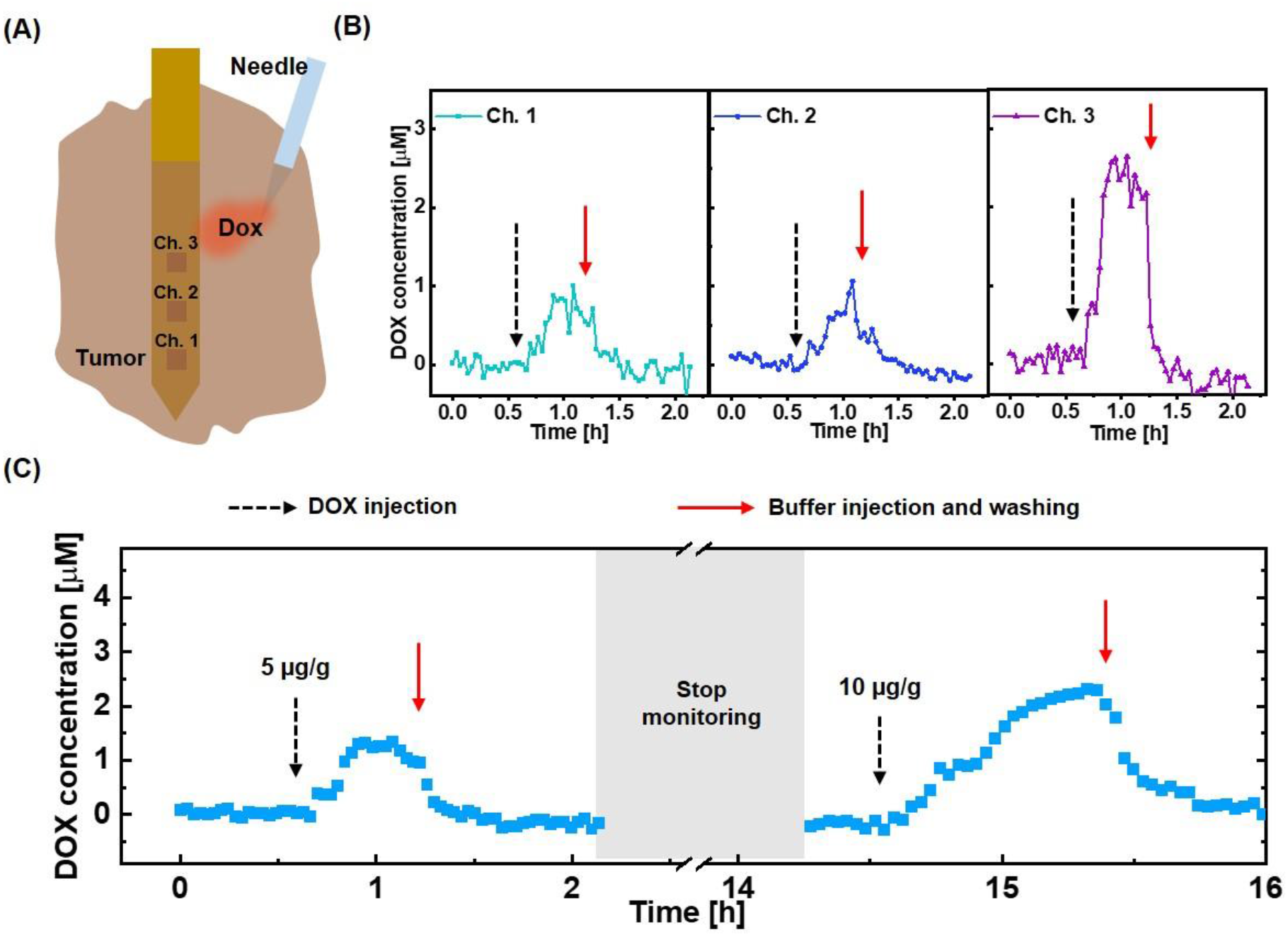
*Ex vivo* monitoring of DOX in tumor tissue. (A) The sensor was implanted into extracted tumor tissue, after which a bolus of DOX was injected. (B) Real-time DOX concentration at each channel after injecting a 5 μg/g bolus of DOX (black dotted arrow) at a tumor site near channel 3, followed by washing with 1x SSC buffer (red solid arrow). (C) Averaged real-time DOX concentrations from the three sensor channels in tumor tissue over the course of 16 h after spiking in boluses of 5 μg/g or 10 μg/g DOX at different time-points. SWV frequency = 200 Hz, amplitude = 50 mV, n = 3 channels.

We were also able to achieve continuous detection for extended periods of time within tumor tissue. We injected boluses of DOX at a tumor site adjacent to the sensor at different time-points and monitored the DOX concentration over the course of 16 h (**Figure 4C**). After applying the first 5 μg/g bolus, we recorded a peak DOX concentration of 1.3 μM, which returned to zero after washing the tumor tissue with SSC. After signal stabilization, we stopped monitoring for 12 h but left the device implanted; when we restarted monitoring, the signal remained at baseline. We subsequently observed a peak DOX concentration of 2.3 μM after injecting a second bolus of 10 μg/g DOX at the same site in the tumor. These results demonstrate that our sensor system can achieve robust and stable drug detection even after extended implantation.

### *In vivo* real-time monitoring of DOX within tumor tissue

Finally, we demonstrated the capability to continuously measure drug PK within tumor tissue in a live mouse. We anesthetized a C57BL/6 mouse, which had a 12-mm-diameter B16-F10 tumor, and implanted our sensor such that all three channels were within the tumor tissue, with channel 3 closest to the surface and channel 1 deepest within the tumor tissue (**Figure S9**). After measuring baseline signal within the tumor tissue, we injected a 10 μg/g bolus of DOX adjacent to channel 1 and observed the response at all three channels to assess its *in vivo* real-time recording capabilities (**Figure S10**). Channel 1 and 2 responded quickly to this DOX injection, while channel 3 did not respond as it was too far from the injection site for measurable quantities of the drug to diffuse.

We next anesthetized a second C57BL/6 mouse, which also had a 12-mm-diameter B16-F10 tumor, and implanted another sensor in the same manner described above. We collected blood from the tail vein using a heparinized capillary tube and inserted an additional third sensor inside the tube. After measuring the baseline signal within the tumor tissue and circulation, we intravenously (i.v.) injected 10 μg/g DOX through the tail vein. We observed different DOX concentration profiles from each of the three channels in the intratumoral probe (**Figure 5A**), indicating that the drug PK exhibits considerable site-dependent variability due to factors including irregular microvasculature density and interstitial fluid pressure across the whole of the tumor tissue.^15, 44–46^ In parallel, we collected blood samples from the tail vein at 0, 5, 30, 60, and 120 min time-points after the i.v. injection to measure circulatory PK. We derived the PK parameters by fitting to a two-compartment model, which describes the changes in drug concentrations in the central (*i.e*. systemic circulation) and peripheral (*i.e*. tumor tissue) compartments (**Table 1**).^25, 47, 48^ From the perspective of the circulation, this model describes the rate of transport of the drug into the tumor (K_d_blood_) as well as the subsequent rate of drug elimination (K_el_) from the body. For the tumor tissue, we derived the PK parameters by fitting to an one-compartment model, which describes the rate of entry of the drug into a particular region of the tumor from the blood (K_a_tumor_) as well as the rate of drug elimination (K_el_) from the tumor.

**Figure 5.**
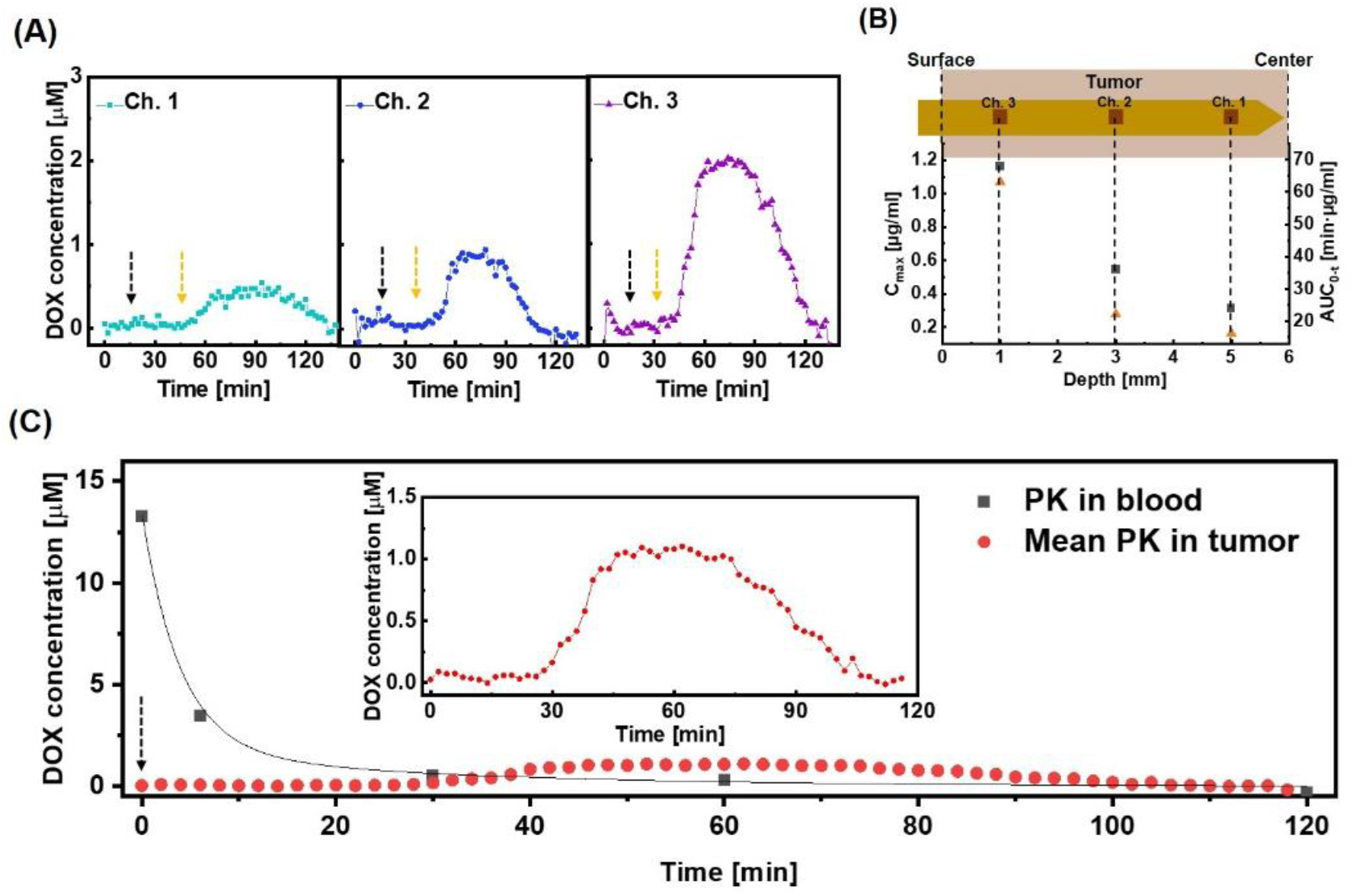
*In vivo* real-time monitoring of DOX PK in tumor tissue. (A) Real-time DOX concentration at each sensor channel after intravenous (i.v.) injection of 10 μg/g DOX at t = 15 min (black dotted arrow). Yellow dotted arrows represent the time-points that the response is observed. (B) Maximum drug concentration (C_max_) (black square) and area under the curve (AUC_0-t_) (orange triangle) as a function of depth within the tumor. (C) Averaged real-time DOX concentrations from the three sensor channels versus DOX concentrations obtained from blood after injecting 10 μg/g DOX at t = 0 min (black dotted arrow). Line represents curve fitting to a two-compartment pharmacokinetic model. Inset is magnified averaged real-time intratumoral DOX concentration data. For (A) and (C), SWV frequency = 200 Hz, amplitude = 50 mV.

**Table 1.** Pharmacokinetic parameters in tumor and blood.

| PK parameter** | In blood | In tumor |  |  |  |
| --- | --- | --- | --- | --- | --- |
|  |  | Ch. 1 | Ch. 2 | Ch. 3 | Mean PK |
| AUC<sub>0-t</sub> [min·μg/ml] | 80.915 | 16.159 | 22.147 | 63.057 | 33.376 |
| C<sub>max</sub> [μg/ml] | 7.702 | 0.312 | 0.546 | 1.163 | 0.625 |
| T<sub>max</sub> [min] | 0 | 79 | 63 | 59 | 64 |
| t<sub>1/2</sub> [min] | 3.4 | 101 | 82 | 90 | 89 |
| K<sub>el</sub> [1/min] | 0.138 ± 0.08 | 0.041 ± 0.005 | 0.067 ± 0.007 | 0.04 ± 0.004 | 0.039 ± 0.004 |
| K<sub>a_tumor</sub> [1/min] | N/A | 0.05 ± 0.007 | 0.077 ± 0.009 | 0.095 ± 0.009 | 0.076 ± 0.009 |
| K<sub>d_blood</sub> [1/min] | 0.064 ± 0.094 | N/A | N/A | N/A | N/A |

\*\*AUC<sub>0-t</sub> is the area under the curve until the drug concentration reaches zero; C<sub>max</sub> is the maximum drug concentration; T<sub>max</sub> is the time at C<sub>max</sub>; t<sub>1/2</sub> is the elimination half-life; K<sub>el</sub> is the elimination rate of drug; K<sub>a\_tumor</sub> is the absorption rate of the drug within the tumor; and K<sub>d\_blood</sub> is the distribution rate of the drug from the bloodstream to the tissue.

Channel 1 was nearest the center of the tumor, where the high interstitial fluid pressure due to the dense microvasculature results in a lower rate of drug penetration relative to the surface of tumor tissue.^49, 50^ Accordingly, we began to observe DOX signal in channel 1 30 min after i.v. injection, followed by a relatively slow increase in drug concentration (K_a_tumor_ = 0.05 min^−1^). We observed a much earlier response in channels 2 and 3—18 min and 15 min after i.v. injection, respectively—and much higher K_a_tumor_ (0.077 min^−1^ and 0.095 min^−1^, respectively).

We observed a clear gradient of decreasing drug exposure as we looked deeper into the tumor (**Figure 5B**). For example, the maximum drug concentration (C_max_) for channel 1 (0.312 μg/ml), which is located at a 5-mm depth within the tumor, was four times lower than for channel 3 (1.163 μg/ml), which is positioned at a 1-mm depth. Similarly, the area under the curve (AUC_0-t_), which reflects the total drug exposure over time, gradually decreased from 63.057 min·μg/ml to 16.159 min·μg/ml as we sampled deeper within the tumor, which would presumably lead to the differential therapeutic efficacy of chemotherapy across the tumor. In parallel, each channel reached C_max_ at an increasingly later time-point post-administration (T_max_) as we looked deeper and deeper into the tumor (**Figure S11**). Interestingly, we noted that the PK parameters did not always fall along a clear continuum from channel 1 to 2 to 3. For example, we measured a lower elimination half-life (t_1/2_) at channel 2 (82 min) than channel 3 (90 min), even though the drug concentration at channel 2 reached C_max_ later than channel 3 (T_max_ = 63 min versus 59 min). We posit that this is due to the fact that the DOX elimination rate (K_el_) was higher in the tissue surrounding channel 2 (0.067 min^−1^) compared to at channel 3 (0.04 min^−1^). These results reveal how the PK can vary across the tumor tissue in a manner that would be difficult or impossible to measure accurately based on conventional biopsy methods.

Finally, we showed that the PK measurements obtained from the tumor tissue differ markedly from the systemic PK (**Figure 5C** and **Table 1**). We made this comparison by determining the mean values for the various PK parameters within the tumor tissue by fitting the average real-time DOX concentrations at all three channels over the course of the experiment. We observed that the total drug exposure (AUC_0-t_) in blood (80.915 min·μg/ml) was ~2.5-fold greater than in the tumor (33.376 min·μg/ml), indicating that the tumor received only a fraction of the drug circulating in the blood. Likewise, C_max_ in blood (7.702 μg/ml) was over ten-fold greater than in tumor tissue (0.625 μg/ml), which raises the possibility of sub-optimal drug dosing if one was to rely entirely on PK data from the bloodstream. The t_1/2_ and K_el_ values were respectively very short (3.4 min) and fast (0.138 min^−1^) in the blood, reflecting rapid clearance by renal excretion.^51^ In contrast, elimination was considerably slower within the tumor (t_1/2_ = 89 min, K_el_ = 0.039 min^−1^). This indicates that the drug remains present and active within this tissue for a longer duration, which could result in an overdose if one were to rely solely on PK measurements from the blood. Conversely, this slower elimination rate could result in a more potent effect even from a seemingly low dose, which is a useful consideration when attempting to identify the appropriate therapeutic window. This striking variability in PK parameters between blood and tumor demonstrates the importance of obtaining local measurements of drug absorption within tumor tissue, and shows how *in situ* tumor measurements collected with our sensor could facilitate the selection of more effective and appropriate drug dosing strategies.

## Conclusion

Drug PK can vary considerably within cancerous tissue relative to the systemic circulation due to the abnormal physiology of the tumor microenvironment. As a consequence, blood-based measurements of drug absorption and elimination are likely to produce misleading or inaccurate PK measurements that impede efforts to identify an optimally safe and effective dose for cancer therapeutics. In this work, we present a microelectrode array-based implantable electrochemical aptamer sensor that overcomes this problem by enabling simultaneous multi-site drug concentration monitoring within tumor tissue in real-time. Our sensor incorporates gold nanoporous electrodes that are highly resistant to both damage and biofouling during or after tissue implantation, and is built on a flexible polymer substrate that matches the physical properties of the surrounding tissue and thus minimizes tissue disruption at the site of insertion. Most importantly, each probe contains three distinct microelectrode channels, and we demonstrate the capacity to sensitively discriminate local differences in the intratumoral concentration of DOX at each channel. Our sensor enabled us to collect extensive tumor-specific PK data over the course of multiple hours that reveal striking differences in the profile of DOX absorption and elimination relative to PK measurements based on systemic circulation from the same animal.

Based on these findings, we believe that our sensor platform could offer a highly effective tool for preclinical analysis of the PK characteristics of experimental drugs, thereby guiding dose selection for first-in-human studies that maximize likelihood of efficacy while minimizing dose-related toxicity. The foundational sensor design demonstrated here should be readily extensible to include larger numbers of channels that can produce measurements with even greater spatial resolution. In principle, different microelectrode channels could also be functionalized with different aptamers, enabling the real-time monitoring of multiple drug agents in the context of a combination therapy, or simultaneous measurement of a therapeutic agent and a biomarker related to drug response. With further refinement and demonstration of the long term stability, safety, and biocompatibility of this probe design, we could envision future adaptations of this platform for potential use in clinical drug and biomarker studies.

## METHODS

### Reagents and materials

The DOX aptamer was obtained from Biosearch Technologies: 5’-HS-C6-ACCATCTGTGTAAGGGGTAAGGGGTGGT-MB-3’, where MB indicates the methylene blue redox reporter. Tris(2-carboxyethyl)phosphine (TCEP), doxorubicin (DOX), and 6-mercapto-1-hexanol (6-MCH) were purchased from Sigma-Aldrich. A diluted 1X saline-sodium citrate (SSC) buffer was prepared by diluting 20X SSC buffer (Thermo Fisher) with nuclease-free water. Fetal bovine serum (FBS) was obtained from Thermo Fisher. Solutions of various DOX concentrations were prepared by dissolving DOX in either 1X SSC buffer or undiluted FBS. The conventional Ag/AgCl reference electrode was prepared from 500-μm-diameter Ag wire (41390 Silver wire, Alfa Aesar) treated with 1 M iron(III) chloride (Sigma-Aldrich) solution for 1 min.

### Fabrication of microelectrode array sensor

A schematic of the device fabrication process is shown in **Figure S2**. For the planar gold microelectrodes, a 300-nm-thick aluminum (Al) sacrificial layer was deposited on a Si wafer by electron-beam evaporation (ATC-E, AJA International Inc.). A 15-μm-thick polyimide (PI) layer (PI2574, HD MicroSystems) was spin-coated onto the Al layer by manual resist spinner (Headway Research) and thermally imidizated by baking in a N_2_-purged oven at 250 °C for 2 h and subsequently cooling down to room temperature RT for 4 h. Then, a 2-μm-thick negative photoresist (PR) layer (NR9-3000PY, Futurrex) was spin-coated onto the PI layer, and then patterned by photolithography with a contact mask aligner (Karl Suss MA-6, SUSS MicroTec) to make the pattern of a 100 x 100 μm^2^ three-channel array. Subsequently, a Ti/Au (5 nm/70 nm) layer was deposited onto the PR layer using electron-beam evaporation, after which the gold planar microelectrode array was formed through a lift-off process in which the PR layer was dissolved in acetone.

For the nanoporous microelectrode arrays, a 4.5-μm-thick positive PR layer (MEGAPOSIT SPR220-3, Kayaku Advanced Materials) was subsequently spin-coated onto the sample. This was patterned to make the pattern of a 100 x 100 μm^2^ three-channel array as described above. A Ti/Au (10 nm/50 nm) bottom-protective layer (BPL) was then deposited onto the PR via sputter deposition (LAB Line SPUTTER, Kurt J. Lesker Co.). A 300-nm-thick Au:Ag alloy layer was then deposited by co-sputtering Au and Ag. The alloy was composed of 66.7% Ag and 33.3% Au. After deposition, the sample was immersed in 69% v/v nitric acid for 7 min at RT to dissolve the Ag, forming a gold nanoporous layer. A lift-off process was then used to form the final gold nanoporous microelectrode array.

For both sensor designs, a 2-μm-thick SU8 encapsulation layer (SU8-2002, Microchem) was spin-coated onto the sample, and then patterned by photolithography with the contact mask aligner to encapsulate the entire sensor with the exception of the microelectrode array. This assembly was then dry-etched with an ICP-RIE etcher (Versaline LL ICP, Plasma-Therm) and Al etching mask to define the shape of the sensor. After removal of the Al etching mask, the sensor was released from the wafer through anodic dissolution of the Al sacrificial layer. For this step, the Si wafer was connected to a DC power supply (1666, B&K Precision), immersed in 2 M NaCl, and 15 V DC voltage was applied to the Al layer.

### Functionalization of aptamer onto the electrode

The DOX aptamer was dissolved in nuclease-free water with a concentration of 100 μM. This solution was reacted with a 1,000-fold molar excess of TCEP solution with a 1:1 volume ratio for 1 h, leading to reduction of the MB moiety and thiol-end group on the aptamer. Afterwards, the freshly-prepared sensor was rinsed with DI water, and then functionalized with 1 μM of the TCEP-treated DOX aptamer in 1x SSC buffer for 2 h at RT. The sensor was then washed with excess buffer and incubated with 7 mM 6-MCH solution for 24 h at RT to passivate the remaining electrode surface. The sensor was stored in 1x SSC at 4 °C until it was used for electrochemical measurement.

### Electrochemical signal measurement

All *in vitro* measurements were performed in a PDMS chamber with the probe connected to a potentiostat (PalmSens4, PalmSens) and multiplexer (MUX8-R2, PalmSens). The PDMS chamber was made by punching a 6-mm-diameter hole on the 5-mm-thick PDMS film and subsequently putting the PDMS film on a glass slide. We used two types of PCB, one small and one large. Each was soldered with an 8-pin FPC connector (FH19C-8S-0.5SH, Hirose Electric Co.). The other side of the large PCB was soldered to another connector (NPD-FF, Omnetics Connector Corporation). The sensor was connected to the small PCB, which was in turn connected to the large PCB. A wire connector (NSD-WD, Omnetics Connector Corporation) connected to the large PCB linked the three channels on the sensor to the commercial potentiostat. We placed the sensor and Ag/AgCl reference electrode into the PDMS chamber and adjusted the height to completely submerges all three channels while avoiding direct contact between the connector and the buffer solution. The PDMS chamber was filled with 1x SSC buffer. Square wave voltammetry (SWV) measurement was carried out over the potential range of −0.55 V to −0.1 V with an amplitude of 50 mV, step size of 1 mV, and pulse frequencies of 200 Hz. Data processing and visualization were performed with custom MATLAB code. All modules were connected as shown in **Figure S4**.

Continuous *in vitro* drug monitoring experiments and biofouling tests in undiluted FBS were conducted in a flowing system that switched between FBS only and FBS plus DOX. Flow was achieved by connecting the PDMS chamber with a peristaltic pump, as shown in **Figure S6**. The inlet and outlet were connected through fluorinated ethylene propylene tubes (Tygon tubing, 1.58 mm inside diameter), which were then mounted onto the peristaltic pump, with a flow-rate of 100 μl/min.

### Melanoma model

All animal studies were performing in accordance with Stanford’s IACUC guidelines and protocols (APLAC protocol # 32947). Murine B16F10 melanoma cells were purchased from ATCC, tested for mycoplasma contamination using the MycoAlert Microplasma Kit (Lonza), and cultured with 0.2-micron filtered DMEM media (Thermo Fisher) supplemented with 10% FBS (Novus Biologicals) and 1% penicillin-streptomycin (Thermo Fisher). C57BL/6 mice (7-8 weeks old; Charles River Laboratories) were subsequently subcutaneously injected with 100 μL of 3 x 10^6^ B16F10 cells/mL in PBS above the right hind leg. Following tumor inoculation, mice were monitored for the formation of palpable tumors, which occurred within 7-10 days. Tumors were regularly monitored via caliper (Mitotoyu) measurements until they reached the appropriate size for either *ex vivo* or *in vivo* studies. Mice were euthanized (CO2 asphyxiation followed by cervical dislocation) when overall tumor area (L x W) exceeded 150 mm^2^, or if mice displayed signs of morbidity (*e.g*., pain, hunching, ulceration, wasting).

### *Ex vivo* tumor experiments

Tumors were surgically removed once they grew to ~100 mm^2^, after euthanasia of the mice. Sensor insertion was tested by inserting the sensor into random positions in the tumor, after which the sensor was immediately withdrawn—this constituted one cycle of insertion. We conducted multiple insertion cycles, and SWV measurements were carried out in 1x SCC buffer after each cycle to monitor sensor function.

DOX detection in *ex vivo* tissues was assessed by placing excised tumors into the PDMS chamber in a small volume of 1x SSC buffer. The sensor was gently implanted into the middle of the tumor tissue to make sure all three channels were inside the tissue. SWV measurement was carried out over the potential range of −0.55 V to −0.1 V with an amplitude of 50 mV, step size of 1 mV, and pulse frequencies of 200 Hz as we injected boluses of DOX into the tumor. Once the measurement was complete, we left the probe in place but washed out the drug by injecting 500 μl of 1x SSC buffer three times into the tumor and collected the waste solution from the PDMS chamber. We then added fresh 1x SSC buffer to the chamber. After signal stabilization, we stopped SWV measurement but left the experimental set-up in place for 12 h. SWV measurement was then carried out again with a second bolus of DOX and an additional washing process.

### *In vivo* experiments

Tumors were allowed to grow to a sufficient size to accommodate the sensor (~140 mm^2^). Mice were then anesthetized using isoflurane (3% induction, 2% maintenance) and maintained on a heating pad at 35 °C in a Faraday Cage (VistaShield, Gamry Instruments). To prevent uneccesary pain or discomfort, mice were subcutaneously injected with buprenorphine-SR (0.5 mg/kg) and Puralube ophthalmic ointment was applied to the eyes. Analgesics were given 15 min to take effect prior to beginning the procedure. Our sensor and Ag/AgCl reference electrode were vertically implanted into the middle of the tumor tissue, and SWV measurement was performed over the potential range of −0.55 V to −0.1 V with an amplitude of 50 mV, step size of 1 mV, and pulse frequencies of 200 Hz. After stabilization of the baseline signal of the sensor, DOX (10 μg/g) was injected through the tail vein manually using a 28G syringe needle. At the end of the experiments, mice were euthanized.

### PK parameter analysis

The DOX concentration-time curve in blood (**Fig. 5C**) was fitted to a bi-exponential equation indicating the two-compartment model:

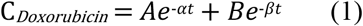

Where A and B are maximum plasma concentrations corresponding to the drug distribution phase and drug elimination phase, respectively, and 1/α and 1/β are the half-lives for distribution and elimination, respectively. C_*Doxorubicin*_ is DOX concentration as a function of time.

K_d_blood_ was calculated by Equation 2:

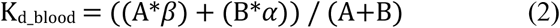

Elimination rate (K_el_) was calculated by Equation 3:

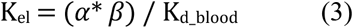

The DOX concentration-time curve within the tumor (**Fig. 5A, C**) were fitted to an exponential equation indicating the one-compartment model:

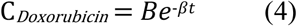

Where β is K_el_ of drug within tumor.

K_a_tumor_ was derived by fitting the absorption phase of the DOX concentration-time curve to Equation 5:

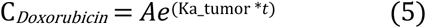

The absorption phase is the time range from drug injection to when the drug concentration reaches C_max_. C_max_, T_max_, t_1/2_ were directly obtained from the experimental raw data of the DOX concentration-time curve. The area under the curve (AUC) was calculated using the definite integral of the DOX concentration-time curve with a time range from drug injection to when the drug concentration reached zero.

## ACKNOWLEDGEMENTS

This work was supported by the Chan-Zuckerberg Biohub, the Biomedical Advanced Research and Development Agency (BARDA, 75A50119C00051) and the National Institutes of Health (NIH, OT2OD025342). Ji Won Seo was supported by Basic Science Research Program through the National Research Foundation of Korea(NRF) funded by the Ministry of Education (2018R1C1B6009140). S.C. is supported by the National Cancer Institute of the National Institutes of Health under Award Number F32CA247352. We thank Celine Liong and Carolyn K. Jons for their thoughtful assistance on the *in vivo* experiments. We thank Leighton Wan, Ian Thompson, and Alyssa Cartwright for their thoughtful comments and valuable suggestions.

## AUTHOR CONTRIBUTION

J.-W.S., K.F., and H.T.S. initiated the project and designed experiments. J.-W.S designed and fabricated the sensor. J.-W.S. conducted experiments and analyzed the data. J.-W.S. and S.C. performed animal experiments. J.-W.S., K.F., S.C., E.A.A., and H.T.S. discussed the data. J.-W.S., K.F., M.E. and H.T.S. wrote the paper. All authors edited the paper.

## Conflict of Interest

The authors declare no conflict of interest.

## Notes

### Competing Interest Statement

The authors have declared no competing interest.

