## Supporting Information for "Real-time monitoring of drug pharmacokinetics within tumor tissue in live animals"

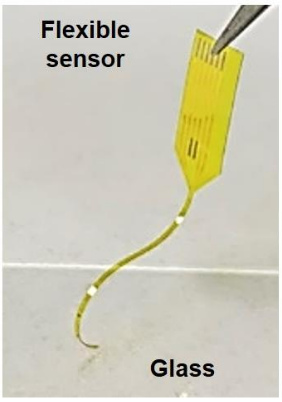


**Figure S1.** Flexible polyimide-based sensor on glass. The probe is shaped to minimize damage at the site of insertion, and offers a good match to the physical properties of surrounding tissue.

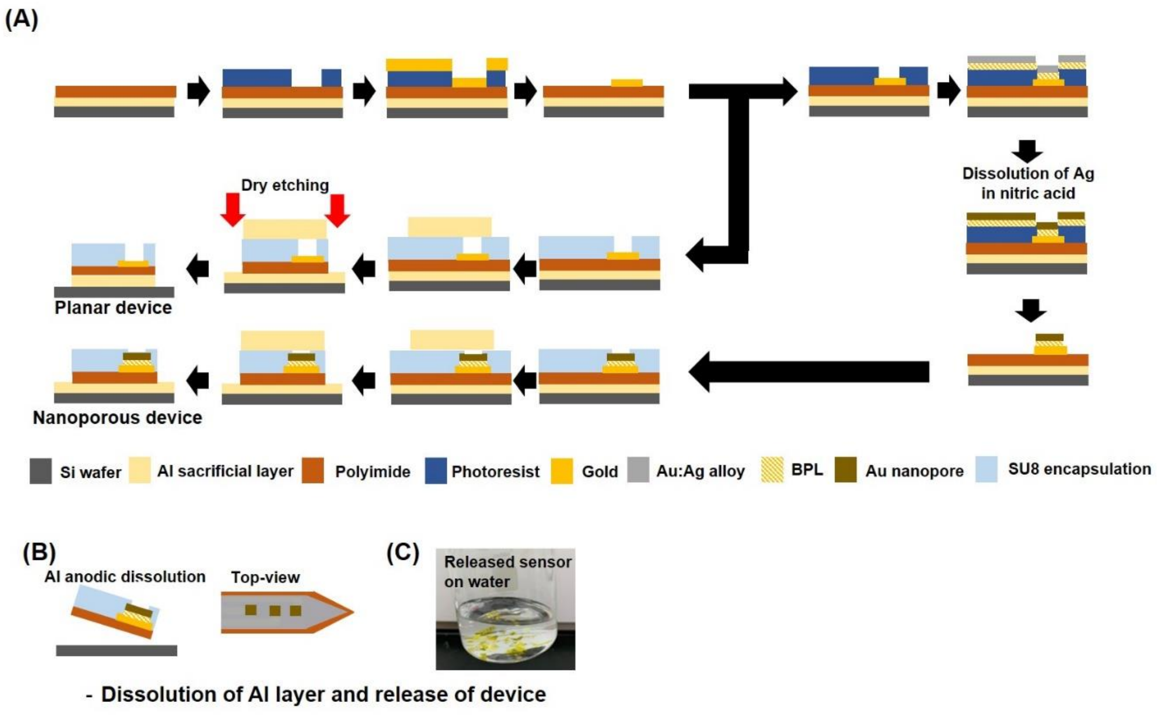


**Figure S2.** Schematic illustration of the fabrication process of gold nanoporous and planar microelectrode array sensor.

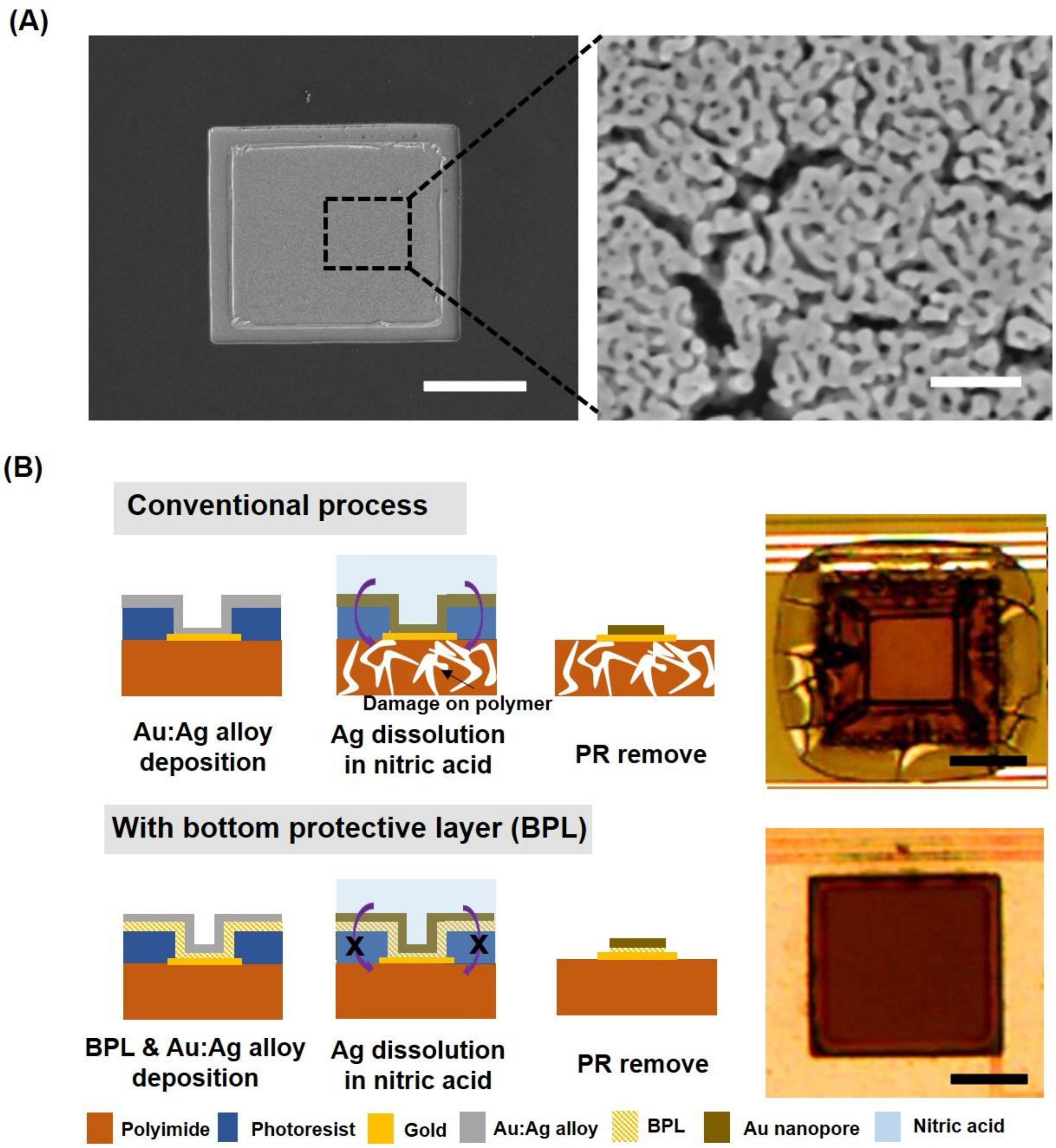


**Figure S3. (**A) SEM image of 100 × 100 μm$^2$ gold nanoporous microelectrodes and magnified image. Scale bar, 50 μm (left) or 50 nm (right). (B) Comparison of the conventional nitric acid dissolution process (top), which degrades the polyimide layer, versus a process that includes a bottom protective layer (BPL) to prevent damage to the polymer (bottom). Righthand panels show corresponding microscopic images of the resulting microelectrodes. Scale bar, 50 μm.

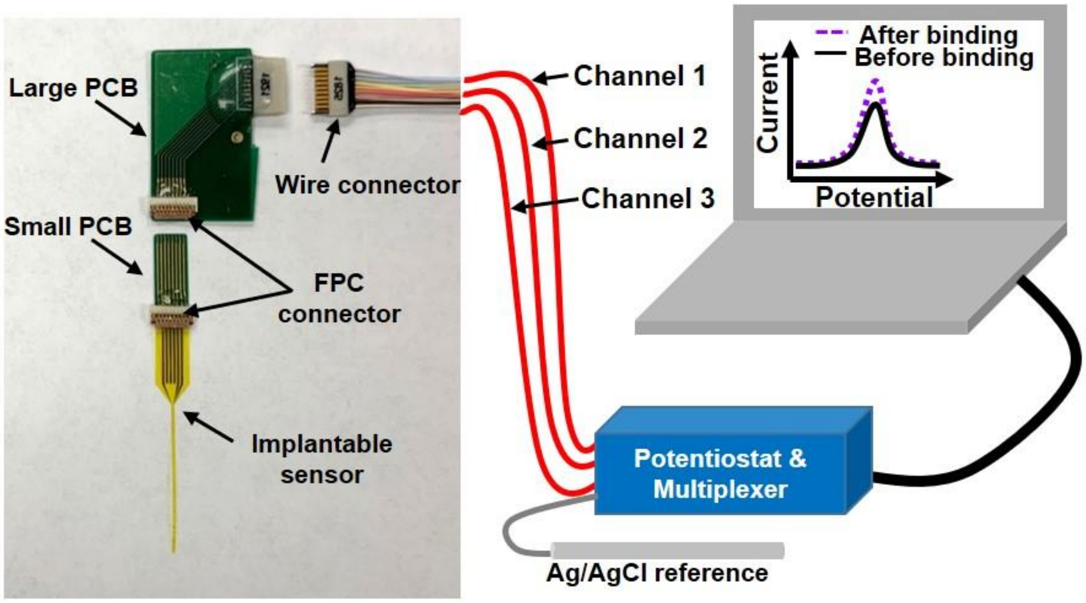


**Figure S4.** Our electrochemical measurement system, including the implantable sensor, FPC connector and printed circuit board (PCB) connection, and commercial potentiostat together with Ag/AgCl reference electrode, connected to a computer with custom Matlab code for real-time data processing and visualization.

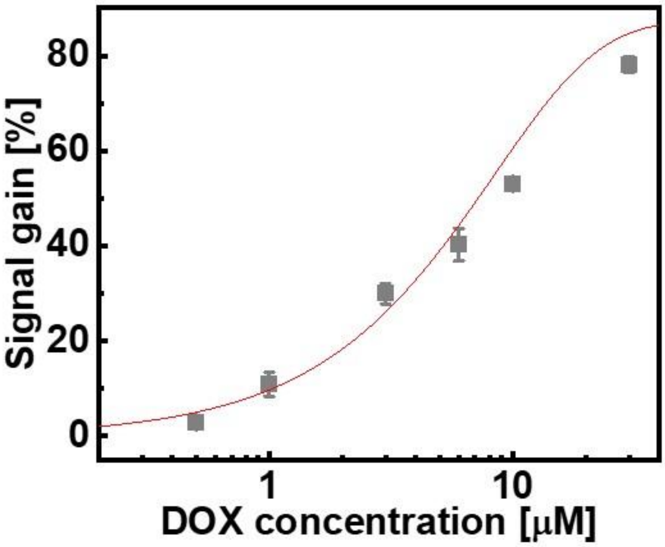


**Figure S5.** Calibration curve fitted to the average signal gain calculated from the multi-channel data shown in Figure 2B. Error bars were calculated from the three-channel data at each concentration.

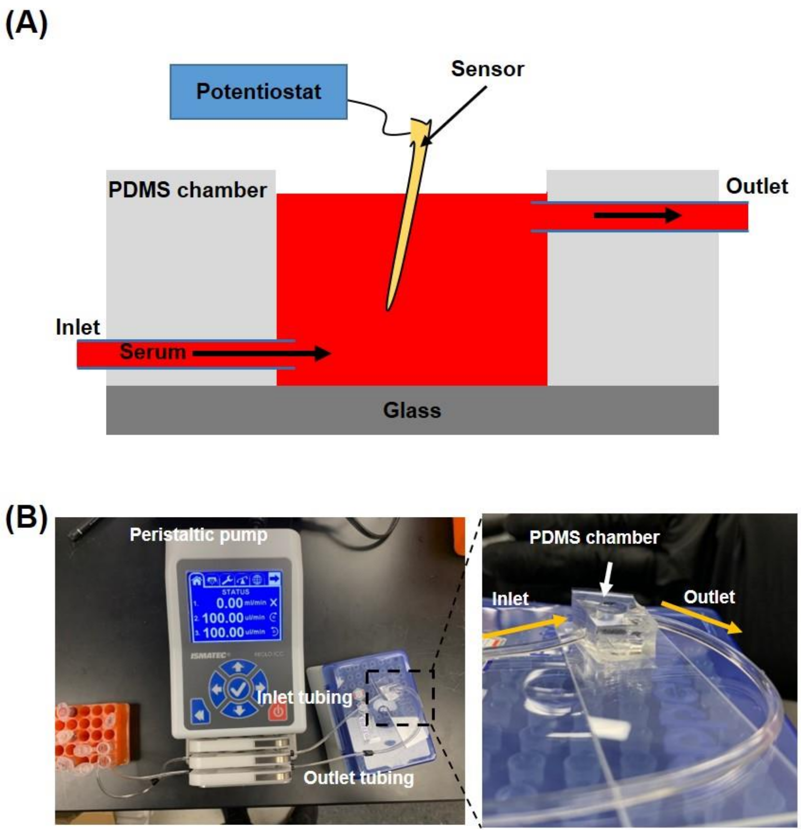


**Figure S6. (**A) Schematic and (B) photo of the system used for testing our sensor in flowing serum.

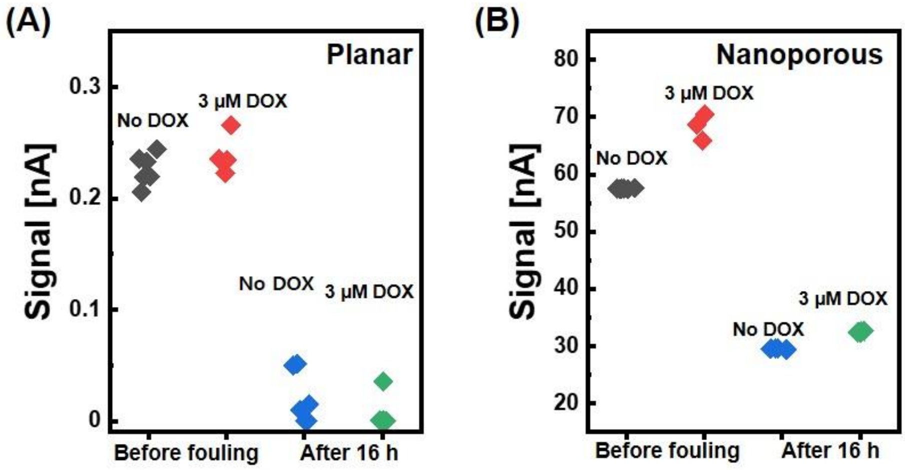


**Figure S7.** Raw signal before and after spiking 3 μM DOX into flowing serum for (A) planar and (B) nanoporous gold microelectrodes. Data are shown for initial signal before biofouling can occur, and for signal obtained after 16 h in blood serum.

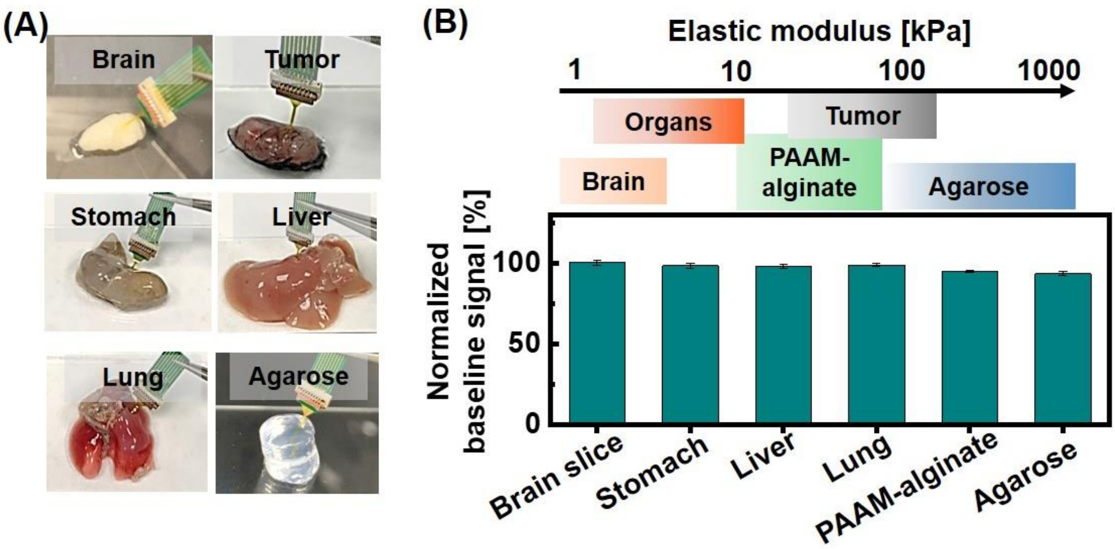


**Figure S8.** (A) Photos of our nanoporous microelectrode sensor inserted into various biological tissues or other materials and (B) the corresponding baseline signal change after 100 insertion cycles. These various materials reflect a broad range of elastic modulus values—the various biological tissues ranged from 1 to 100 kPa, and we also included PAAM-alginate hydrogel (30 kPa) and agarose hydrogel (1,000 kPa).

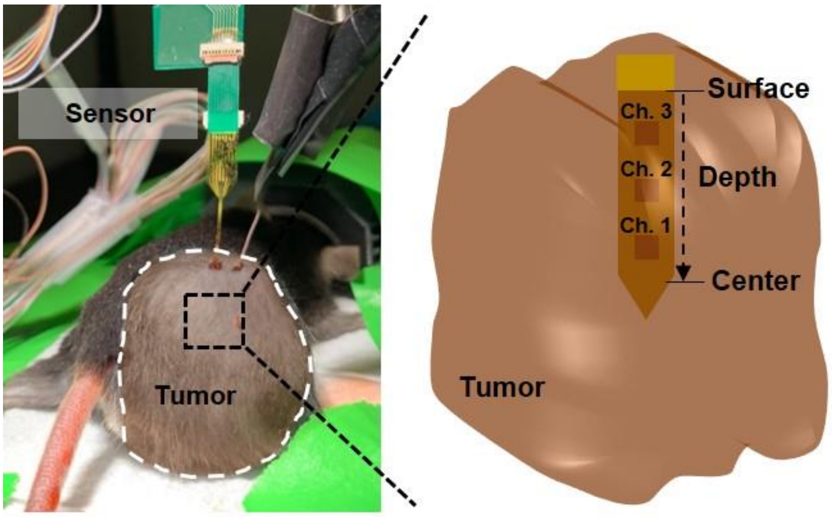


**Figure S9.** Microelectrode array sensor implantation into tumor tissue of an anesthetized mouse.

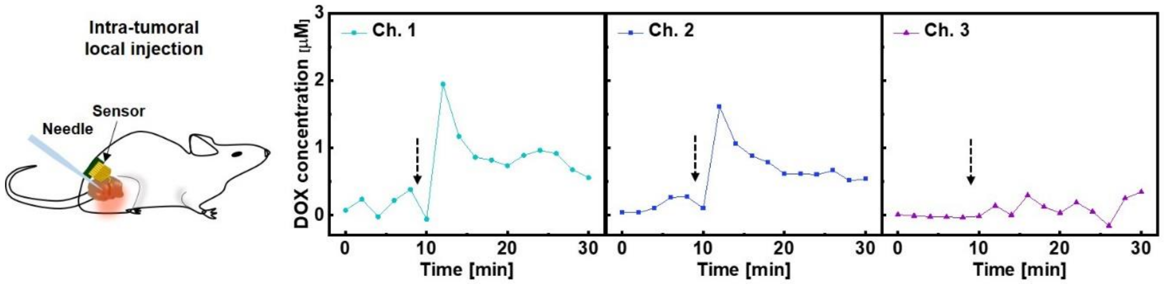


**Figure S10.** Real-time DOX concentration measurements at each sensor channel after intra-tumoral injection of 10 µg/g DOX near channel 1 at t = 9 min (black dotted arrow).

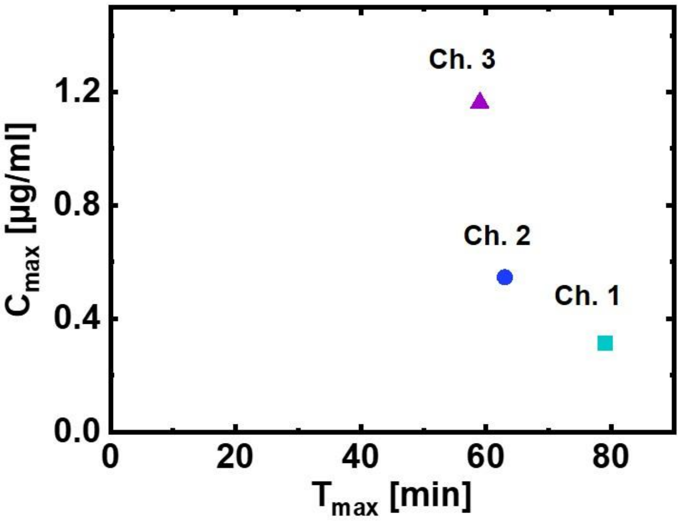


**Figure S11.** Maximum drug concentration ($C_{max}$) and time at $C_{max}$ ($T_{max}$) from the three sensor channels.
